# Evaluation and application of summary statistic imputation to discover new height-associated loci

**DOI:** 10.1101/204560

**Authors:** Sina Rüeger, Aaron McDaid, Zoltán Kutalik

## Abstract

**Abstract:** As most of the heritability of complex traits is attributed to common and low frequency genetic variants, imputing them by combining genotyping chips and large sequenced reference panels is the most cost-effective approach to discover the genetic basis of these traits. Association summary statistics from genome-wide meta-analyses are available for hundreds of traits. Updating these to ever-increasing reference panels is very cumbersome as it requires reimputation of the genetic data, rerunning the association scan, and meta-analysing the results. A much more efficient method is to directly impute the summary statistics, termed as *summary statistics imputation*. Its performance relative to *genotype imputation* and practical utility has not yet been fully investigated. To this end, we compared the two approaches on real (genotyped and imputed) data from 120K samples from the UK Biobank and show that, while *genotype imputation* boasts a 2- to 5-fold lower root-mean-square error, *summary statistics imputation* better distinguishes true associations from null ones: We observed the largest differences in power for variants with low minor allele frequency and low imputation quality. For fixed false positive rates of 0.001, 0.01, 0.05, using *summary statistics imputation* yielded an increase in statistical power by 15, 10 and 3%, respectively. To test its capacity to discover novel associations, we applied *summary statistics imputation* to the GIANT height meta-analysis summary statistics covering HapMap variants, and identified 34 novel loci, 19 of which replicated using data in the UK Biobank. Additionally, we successfully replicated 55 out of the 111 variants published in an exome chip study. Our study demonstrates that *summary statistics imputation* is a very efficient and cost-effective way to identify and fine-map trait-associated loci. Moreover, the ability to impute summary statistics is important for follow-up analyses, such as Mendelian randomisation or LD-score regression.

**Author summary:** Genome-wide association studies (GWASs) quantify the effect of genetic variants and traits, such as height. Such estimates are called *association summary statistics* and are typically publicly shared through publication. Typically, GWASs are carried out by genotyping ~ 500′000 SNVs for each individual which are then combined with sequenced reference panels to infer untyped SNVs in each’ individuals genome. This process of *genotype imputation* is resource intensive and can therefore be a limitation when combining many GWASs. An alternative approach is to bypass the use of individual data and directly impute summary statistics. In our work we compare the performance of *summary statistics imputation* to *genotype imputation*. Although we observe a 2- to 5-fold lower RMSE for *genotype imputation* compared to *summary statistics imputation*, *summary statistics imputation* better distinguishes true associations from null results. Furthermore, we demonstrate the potential of *summary statistics imputation* by presenting 34 novel height-associated loci, 19 of which were confirmed in UK Biobank. Our study demonstrates that given current reference panels, *summary statistics imputation* is a very efficient and cost-effective way to identify common or low-frequency trait-associated loci.

## Introduction

Genome-wide association studies (GWASs) have been successfully applied to reveal genetic markers associated with hundreds of traits and diseases. The genotyping arrays used in these studies only interrogate a small proportion of the genome and are therefore typically unable to pinpoint the causal variant. Such arrays have been designed to be cost-effective and include only a set of tag single nucleotide variants (SNVs) that allow the inference of many other unmeasured markers. To date, thousands of individuals have been sequenced [1,2] to provide high resolution haplotypes for *genotype imputation* tools such as **IMPUTE** and **minimac** [3,4], which are able to infer sequence variants with ever-increasing accuracy as the reference haplotype set grows.

Downstream analyses such as Mendelian randomisation [5], approximate conditional analysis [6], heritability estimation [7], and enrichment analysis using high resolution annotation (such as DHS) [8] often require genome-wide association results at the highest possible genomic resolution. *Summary statistics imputation* has been proposed as a solution that only requires summary statistics and the linkage disequilibrium (LD) information estimated from the latest sequencing panel to directly impute up-to-date meta-analysis summary statistics [9]. In a recent paper [10] we improved the most recent *summary statistics imputation* method [11] by extending imputation to account for variable sample sizes and reference panel composition. While we demonstrated in our paper the benefits of new features compared to other similar methods using simulated data, this study compares it directly to genotype imputation and focuses on the practical advantages of *summary statistics imputation* using real data. In particular, we evaluated two experiments: (1) we ran a GWAS on human height using data from 336′474 individuals from the UK Biobank and compared the performances of *summary statistics imputation* and *genotype imputation*, using direct genotyping/sequencing as gold standard; (2) we imputed association summary statistics from a HapMap-based GWAS study [12] using the UK10K reference panel to explore new potential height-associated variants which we validated using results from Marouli *et al.* [13] and the UK Biobank height GWAS.

### Summary statistics imputation

By combining summary statistics for a set of variants and the fine-scale LD structure in the same region, we can estimate summary statistics of new, untyped variants at the same locus. We can formally write this [9–11] using the conditional expectation of a multivariate normal distribution.

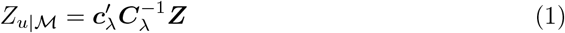

In Equation (1) we aim to impute the Z-statistic of a variant *u*. Vector ***Z*** contains the known Z-statistics of a set of *m* SNVs 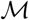, estimated from *N* samples. This set of *m* SNVs are called *tag* SNVs. *C* contains the pairwise correlations among the tag SNVs 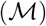, and *c* represents the correlations between 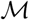 and SNV *u*. Both correlation entities are regularised [14] using a regularisation parameter λ, yielding ***C***_λ_ and ***c***_λ_. In this paper we use 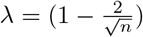 [15]. 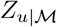 describes the Z-statistic of an untyped SNV *u* given the Z-statistics of a set tag SNVs 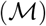.

Since LD between SNVs is minimal beyond 250 Kb, we choose 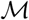 to include all measured variants within at least 250 Kb from SNV *u*. To speed up the computation when imputing SNVs genome-wide, we apply a sliding window strategy, where SNVs within a 1 Mb window are imputed simultaneously using the same set of *m* tag SNVs within the 1 Mb window ± 250 Kb flanking regions.

We use an adjusted imputation quality that corrects for the effective number of tag SNVs *p*_eff_ [16]:

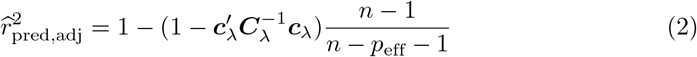

To account for variable sample size in summary statistics of tag SNVs, we use an approach to down-weight entries in the ***C***_λ_ and ***c***_λ_ matrices for which summary statistics were estimated from a GWAS sample size lower than the maximum sample size in that dataset [10].

For more details on our *summary statistics imputation* method and extensions of it, see our complementary paper [10].

## Results

To assess the performance of *summary statistics imputation* in realistic scenarios we used two different datasets. In Section “Comparison of *summary statistics imputation* versus *genotype imputation*” we compare the performance of *summary statistics imputation* to *genotype imputation*, using measured and imputed genotype data from 120′086 individuals in the UK Biobank. In Section “*Summary statistics imputation* of the height GWAS of the GIANT consortium”, we use published association summary statistics from 253′288 individuals to show that *summary statistics imputation* can be used to identify novel associations. Both analyses are centered around the genetics of human height. In the following we will often refer to two GIANT (Genetic Investigation of ANthropometric Traits) publications: Wood *et al*. [12], an analysis of HapMap variants that revealed 423 loci, and Marouli *et al*. [13], an exome chip based analysis that revealed 120 new height-associated loci. Together, these two studies — the HapMap and the exome chip study — constitute the most complete collection of genetic associations with height.

### Comparison of *summary statistics imputation* versus *genotype imputation*

By having two types of genetic data at hand, genotype and imputed genotype data, we were able to compare summary statistics of 37′467 typed SNVs resulting from (1) associations calculated from original genotype data (ground truth); (2) associations calculated from imputed genotype data (*genotype imputation*) and (3) associations imputed from summary statistics calculated using genotype data. Fig 1 gives an overview of how these three types of summary statistics are related and compared. For our analysis, we defined 706 genomic regions in total, among which 535 contain SNVs associated with height [12,13], while the remaining 171 regions were selected to be free of any known height associated SNVs.

**Fig 1.**
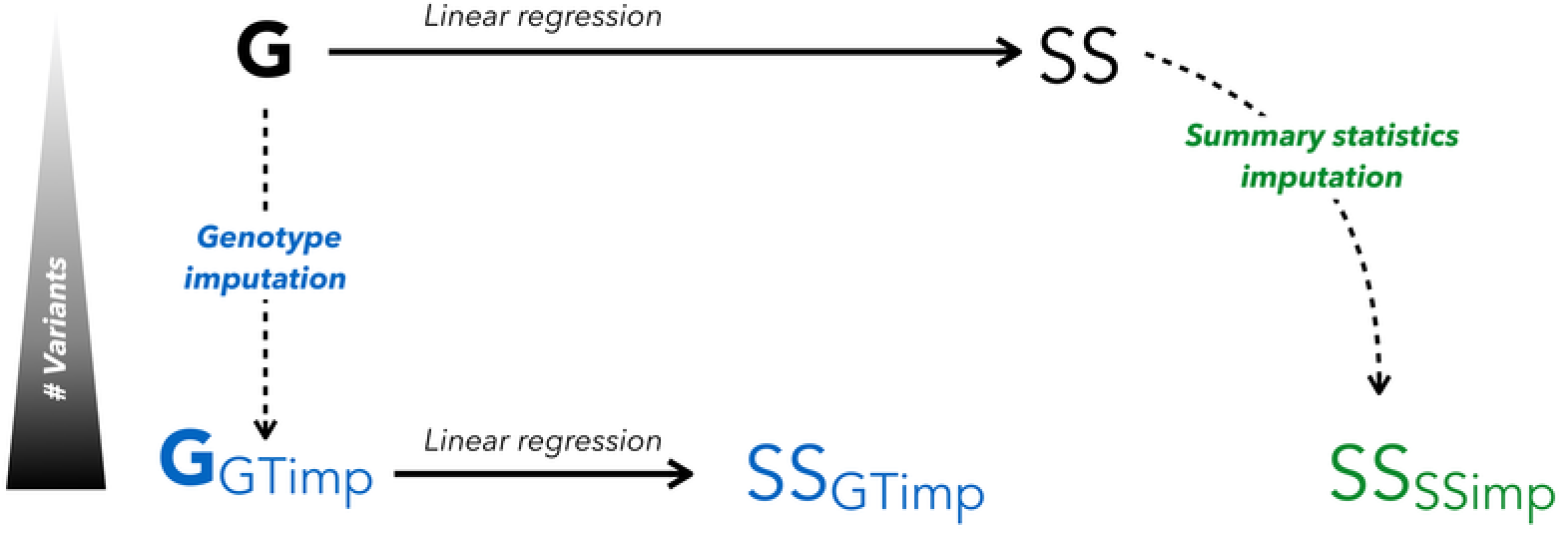
Genotype vs. summary statistics imputation. From genotype data (top-left, **G**) we can calculate summary statistics (top-right, **SS**). Summary statistics for an unmeasured/masked SNV can be obtained via two ways: we can impute genotype data (bottom-left, **G-GTimp**) using *genotype imputation* and then calculate summary statistics via linear regression (bottom-middle, **SS-GTimp**), or by applying *summary statistics imputation* on the summary statistics calculated from genotype data (bottom-right, **SS-SSimp**). For the purpose of our analysis, we are only looking at genotyped (and genotype imputed) SNVs, thus masking one SNV at the time and imputing it using summary statistics from neighbouring SNVs

We examined imputation results for different SNV categories. These were grouped based on (i) their association status (being correlated with the causal SNV *vs*. null SNVs) with the lead SNV of each of the 535 height-associated regions (6′080 variants were correlated, 31′567 were not); (ii) frequency (MAF: 1% < low-frequency ≤ 5% < common; 13′857 and 23′790 variants, respectively); and (iii) imputation quality based on *summary statistics imputation* (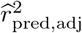: low ≤ 0.3 < medium ≤ 0.7 < high; 10′953, 9′385, and 17′309 variants, respectively). S1 Fig and S2 Fig show the distribution of SNV counts in each of these twelve subgroups. We term the 6′080 SNVs correlated with a height-associated lead SNV as *associated* SNVs. Conversely, we refer to the 31′567 SNVs that are not correlated with any height-associated lead SNV as *null* SNVs. For both, null and associated SNV groups, the largest group of analysed variants were common and well-imputed (S1 Fig). The fraction of SNVs with low quality imputation increases with lower minor allele frequency (S2 Fig). However, the number of rare variants (MAF< 1%) were too small (2′411 variants, among these only 13 associated variants) to draw meaningful conclusions and hence we limited our analysis to common and low-frequency variants.

We focused on two aspects of the imputation results. First, we compared how *summary statistics imputation* and *genotype imputation* perform relative to the ground truth (direct genotyping). For this we used four measures: the root mean squared error (RMSE), bias, the linear regression slope, and the correlation. Second, we calculated power and false positive rate for *genotype imputation* and *summary statistics imputation* directly.

### *Genotype imputation* outperforms *summary statistics imputation* for low allele frequency

Fig 2 shows in green the comparison between summary statistics resulting from measured genotype data (ground truth) and imputed summary statistics for 6′080 height-associated variants. As expected, the performance drops as the imputation quality and as the MAF decrease. For well-imputed common SNVs (the largest subgroup with 4′782 variants), *summary statistics imputation* performs on average well with a correlation and a slope close to 1 (cor = 0.998 and slope = 0.98), but it drops to 0.90 (cor = 0.961 and slope=0.90) for low imputation quality, low-frequency variants. On the other hand, for *genotype imputation* (Fig 2, blue dots) all subgroups of SNVs show near perfect slope and correlation. Note that imputation quality for *summary statistics imputation* and *genotype imputation* differ in definition and we find that the latter was consistently higher (S3 Fig and S4 Fig) and showed little variation across SNVs. To be able to compare the performance between *genotype imputation* and *summary statistics imputation* for the same subgroups of SNVs we used the imputation quality defined by *summary statistics imputation* to classify SNVs.

**Fig 2.**
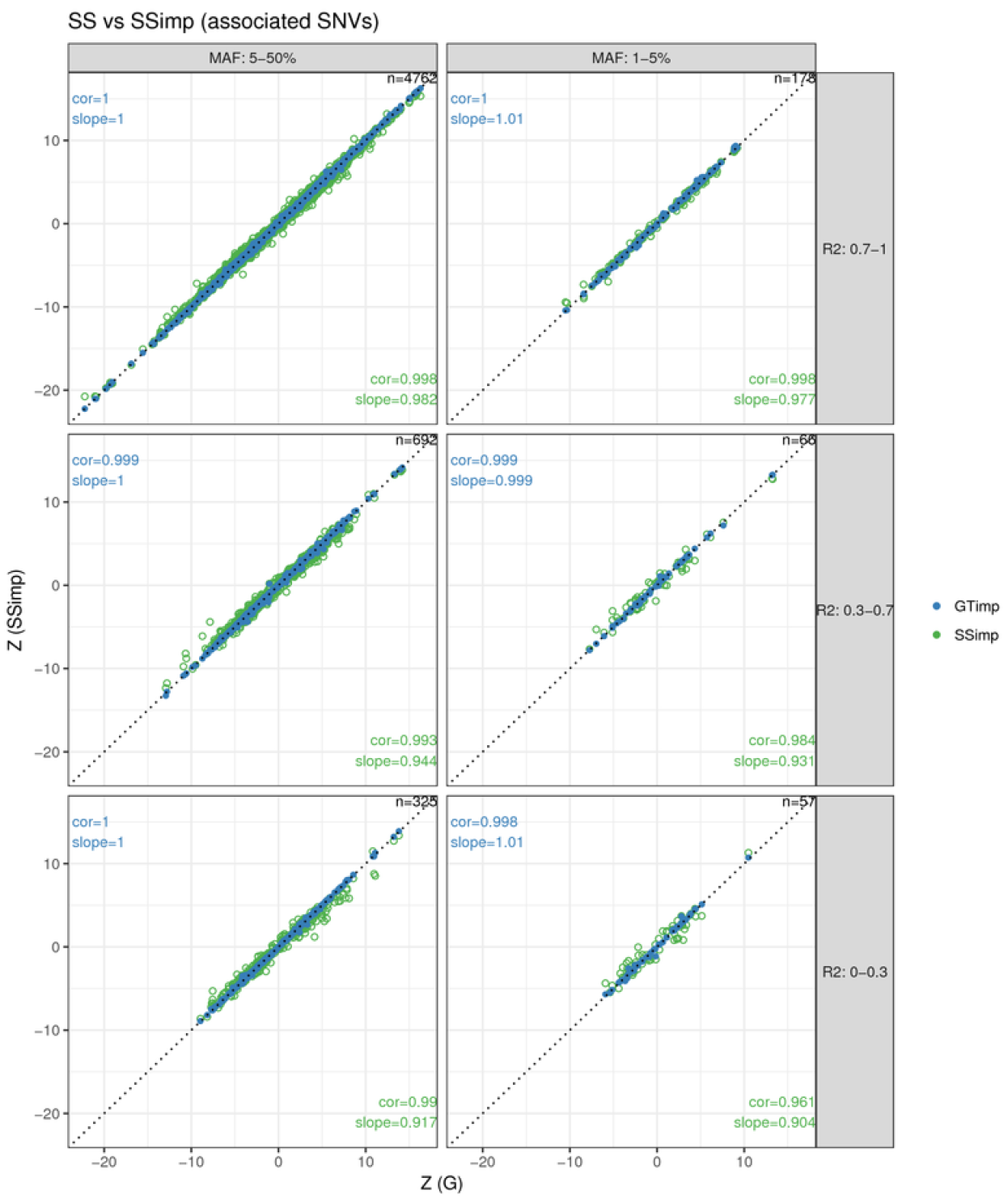
*Summary statistics imputation* versus *genotype imputation* in associated variants. The x-axis shows the Z-statistics of the genotype data (ground truth), while the y-axis shows the Z-statistics from *summary statistics imputation* (green) or *genotype imputation* (blue). Results are grouped according to MAF (columns) and imputation quality (rows) categories and the numbers top-right in each window refers to the number of SNVs represented. The identity line is indicated with a dotted line. The estimation for correlation and slope are noted in the bottom-right corner for *summary statistics imputation* and in the top-left corner for *genotype imputation*. Blue dots are plotted over the green ones.

For the 31′567 null SNVs we present the same metrics as for associated SNVs. We analysed 13′556 low-frequency and 18′011 common variants. First, the green dots in Fig 3 show summary statistics from genotype data and *summary statistics imputation*. We find that both the correlation and slope gradually decrease with dropping imputation quality and MAF. For example, the correlation is 0.91−0.96 for well-imputed, 0.86−0.90 for medium and 0.70−0.82 for badly-imputed SNVs. The blue dots in Fig 3 show the respective results for *genotype imputation*, which exhibits an almost perfect (> 0.98) slope and correlation.

**Fig 3.**
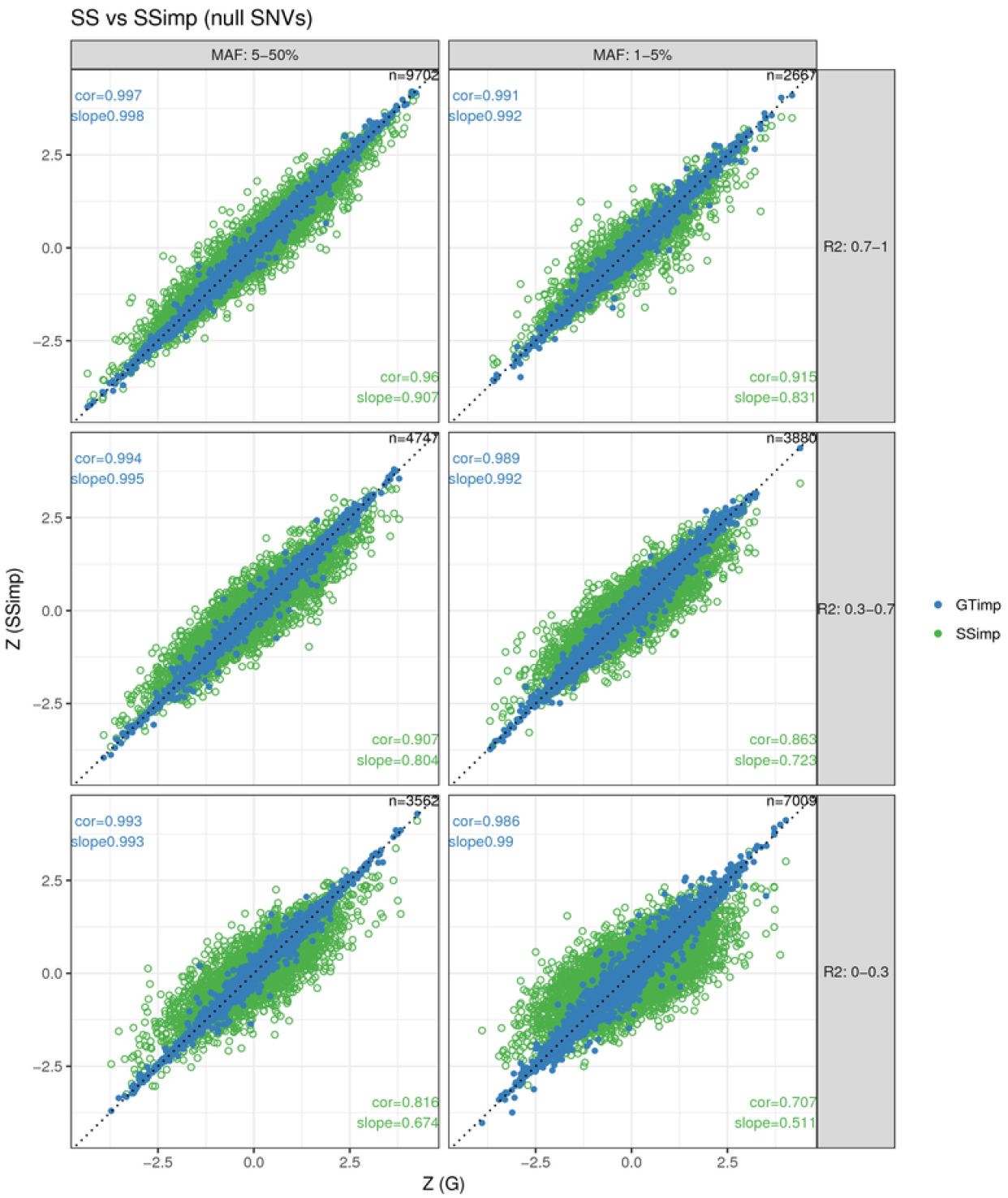
*Summary statistics imputation* versus *genotype imputation* in null variants. The x-axis shows the Z-statistics of the genotype data (ground truth), while the y-axis shows the Z-statistics from *summary statistics imputation* (green) or *genotype imputation* (blue). Results are grouped according to MAF (columns) and imputation quality (rows) categories and the numbers top-right in each window refers to the number of SNVs represented. The identity line is indicated with a dotted line. The estimation for correlation and slope are noted in the bottom-right corner for *summary statistics imputation* and in the top-left corner for *genotype imputation*. Blue dots are plotted over the green ones.

### Effect estimate accuracy and precision

We then compared *summary statistics imputation* and *genotype imputation* in terms of RMSE among associated variants (for the same six SNV categories), shown in the upper part of Table 1. For all six subgroups, *genotype imputation* had a smaller RMSE than *summary statistics imputation*. The difference between the two methods in terms of RMSE increases as imputation quality decreases. For the largest SNV subgroup — well-imputed and common SNVs — *summary statistics imputation* had a RMSE of 0.29 versus 0.085 for *genotype imputation*. In case of *summary statistics imputation*, the RMSE is more influenced by a decrease in imputation quality than by a reduction of MAF. For example, the RMSE for common variants with medium-quality imputation is 0. 51 (a 1.79-increase), while the RMSE for low-frequency variants with high-quality imputation is 0.33 (a 1.16-fold increase). However, for *genotype imputation* a decrease in MAF or imputation quality seems to have a similar effect. For example, the RMSE for well-imputed, low-frequency variants is 0.14 for *genotype imputation* (a 1. 65-increase), and the RMSE for medium-imputed, common variants is 0.13 for *genotype imputation* (a 1.57-increase) (Fig 4). For null SNVs we observe for *summary statistics imputation* a RMSE of 0.33 for well-imputed and common SNVs up to 0.75 for badly-imputed and low-frequency SNVs (lower part in Table 1). For *genotype imputation* the RMSE ranges are much lower, between 0.10 for well-imputed and common SNVs and 0.18 for badly-imputed and low-frequency SNVs. The bias is very close to zero for both approaches and for null and associated SNVs, and does not significantly vary with MAF or imputation quality.

**Table 1.** RMSE for *summary statistics imputation and genotype imputation*.

| | MAF | $\hat{r}_{\text{pred,adj}}^2$ | SSimp<br>RMSE | Bias | GTimp<br>RMSE | Bias | # SNVs |
| --- | --- | --- | --- | --- | --- | --- | --- |
| Associated | 1-5% | 0-0.3 | 0.8979 | 0.0610 | 0.2354 | 0.0197 | 57 |
|  |  | 0.3-0.7 | 0.7195 | 0.0384 | 0.1423 | 0.0076 | 66 |
|  |  | 0.7-1 | 0.3310 | -0.0169 | 0.1408 | 0.0052 | 178 |
|  | 5-50% | 0-0.3 | 0.6366 | -0.0447 | 0.1169 | -0.0018 | 325 |
|  |  | 0.3-0.7 | 0.5135 | 0.0015 | 0.1346 | -0.0102 | 692 |
|  |  | 0.7-1 | 0.2859 | -0.0005 | 0.0854 | -0.0012 | 4762 |
| Null | 1-5% | 0-0.3 | 0.7478 | 0.0020 | 0.1787 | 0.0030 | 7009 |
|  |  | 0.3-0.7 | 0.5431 | -0.0049 | 0.1569 | -0.0001 | 3880 |
|  |  | 0.7-1 | 0.4487 | -0.0034 | 0.1470 | 0.0005 | 2667 |
|  | 5-50% | 0-0.3 | 0.6133 | -0.0122 | 0.1236 | 0.0007 | 3562 |
|  |  | 0.3-0.7 | 0.4788 | -0.0075 | 0.1197 | -0.0048 | 4747 |
|  |  | 0.7-1 | 0.3251 | 0.0038 | 0.0967 | 0.0010 | 9702 |

**Fig 4.**
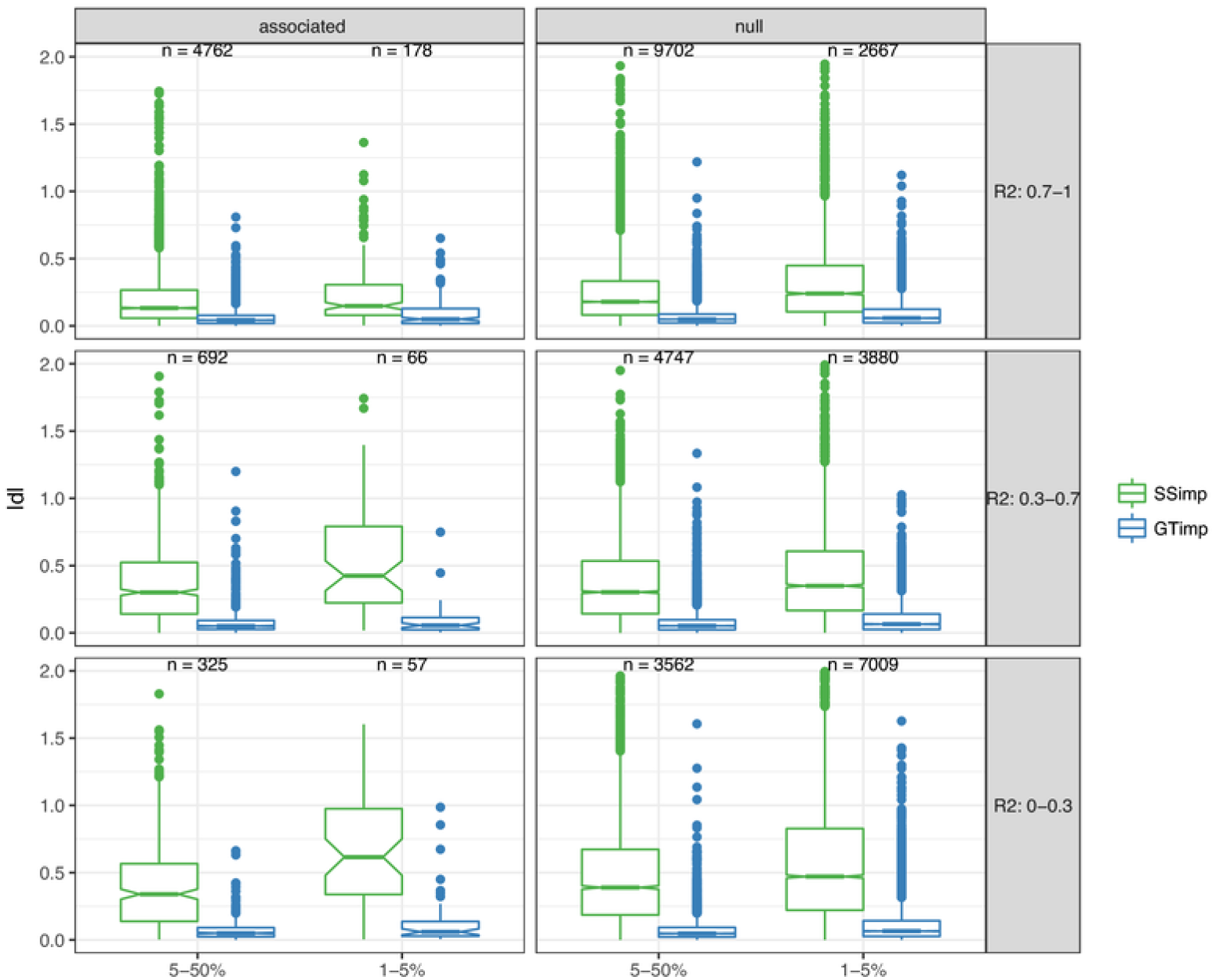
Visualising RMSE of *summary statistics imputation* and *genotype imputation*. This figure uses boxplots to compare the absolute difference |*d*| (used for calculation of RMSE) for each variant between Z-statistics of *summary statistics imputation* (SSimp, green) and *genotype imputation* (GTimp, blue) of associated SNVs (left column) and null SNVs (right column). Results are grouped according to MAF (x-axis) and imputation quality (rows) categories. The numbers printed above the boxplot represents the number of SNVs used for the |*d*| calculation in that MAF and imputation quality subgroup. The corresponding 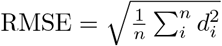 is shown in Table 1.

### *Summary statistics imputation* displays lower false positive rate

Finally, analogous to a ROC curve Fig 5 presents simultaneously power and false positive rate (FPR) with varying significance threshold (*α* from 0 to 1). As before, we stratified the results by MAF and imputation quality categories. We observe that for common SNVs with 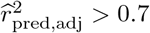 the results for *genotype imputation* and *summary statistics imputation* are differing only by a little with respect to FPR and power. For low-frequency and well-imputed variants, *summary statistics imputation* seems to offer an advantage compared to *genotype imputation* when *α* is low. As we approach lower imputation quality and MAF, *summary statistics imputation* advantage becomes more and more apparent for all range of *α* values. In general, whenever *summary statistics imputation* is outperforming *genotype imputation*, this is because lower FPR (horizontal shift), and not due to increased power. This aspect is more clearly visible in S5 Fig (same data, but untransformed x-axis ranging from 0 to 1). For fixed false positive rates of 0.001, 0.01, 0.05, using *summary statistics imputation* yielded an increase in statistical power by 15, 10 and 3%, respectively.

**Fig 5.**
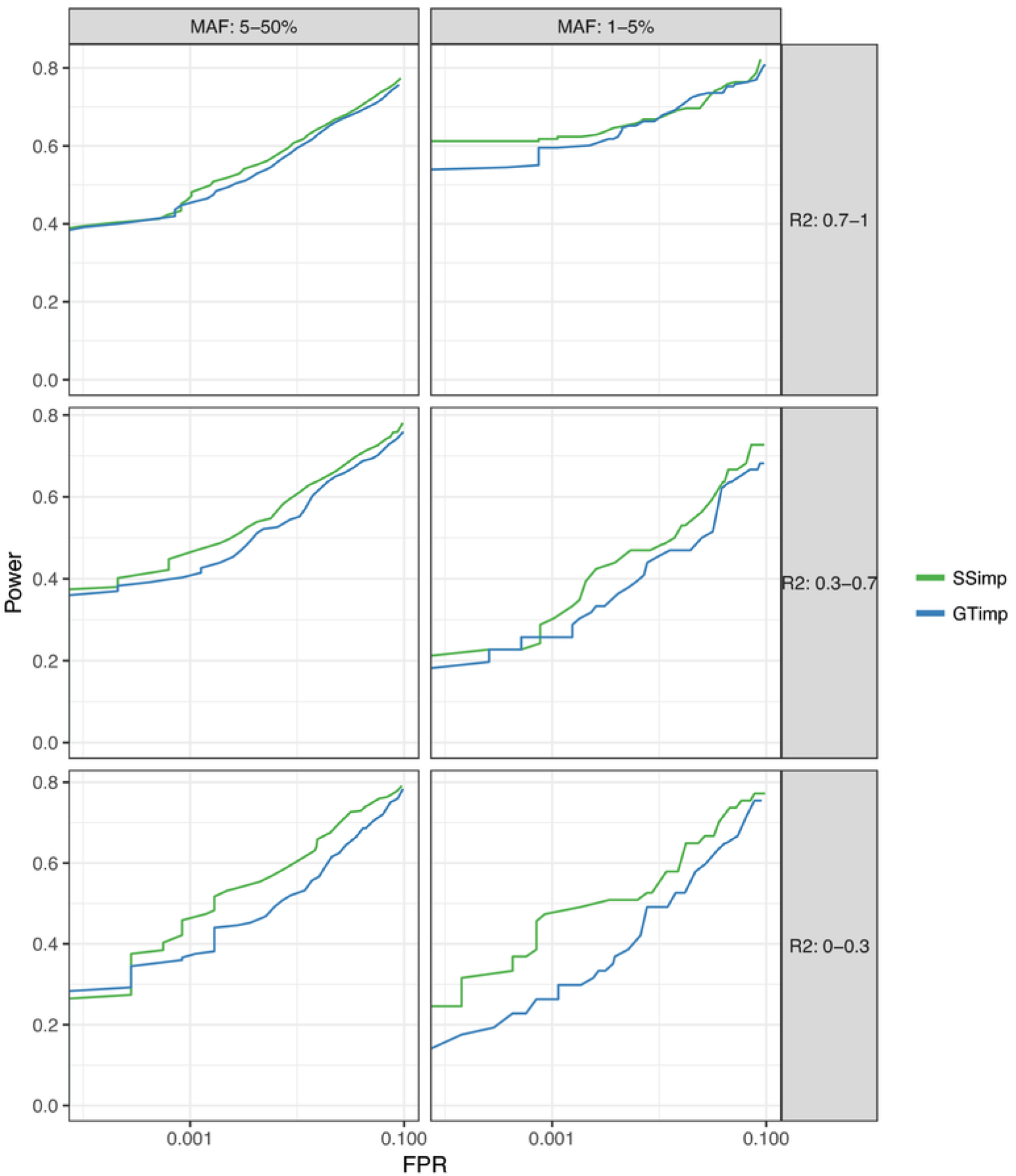
FPR versus power. This figure compares false positive rate (FPR) (x-axis on log10-scale) versus power (y-axis) for *genotype imputation* (blue) and *summary statistics imputation* (green) for different significance thresholds. This figure is a zoom into the bottom-left area of S5 Fig and shows FPR between 0 and 0.1. Results are grouped according to MAF (columns) and imputation quality (rows) categories.

### *Summary statistics imputation* of the height GWAS of the GIANT consortium

While previous studies have examined the role of (common) HapMap variants for height [12,17], the impact of rare coding variants could not be investigated until bespoke genotyping chips (interrogating low-frequency and rare coding variants) were designed to address this question in a cost-effective manner. Such an exome chip based study was conducted by the GIANT consortium in 381′000 individuals and revealed 120 height-associated loci, of which 83 loci were rare or low-frequency [13]. These association results enabled us to compare the usefulness of imputation-based inference with direct genotyping done in Wood *et al*. [12], since the two studies are highly comparable in terms of ancestry composition and statistical analysis, evidenced by S6 Fig confirming very high concordance between summary statistics for the subset of 2′601 SNVs correlated to a height-associated variant which were available in both studies.

### Discovery and replication of 19 new loci

By imputing > 6M additional SNVs summary statistics using HapMap variants [12] as tag SNPs we were interested in two aspects: (1) discovering new height-associated candidate loci, and (2) replicating these candidate loci in the UK Biobank and the GIANT exome chip look-up (Fig 6). We used the HapMap-based height study and the UK10K reference panel as inputs for *summary statistics imputation* and used all HapMap SNVs as tag SNVs. We imputed variants that were available in UK10K with a MAF_UK10K_ ≥ 0.1%, as well as all reported exome variants in Marouli *et al*. [13]. In total we imputed 10′966′111 variants, of which 9′276′018 (84%) had an imputation quality ≥ 0.3.

**Fig 6.**
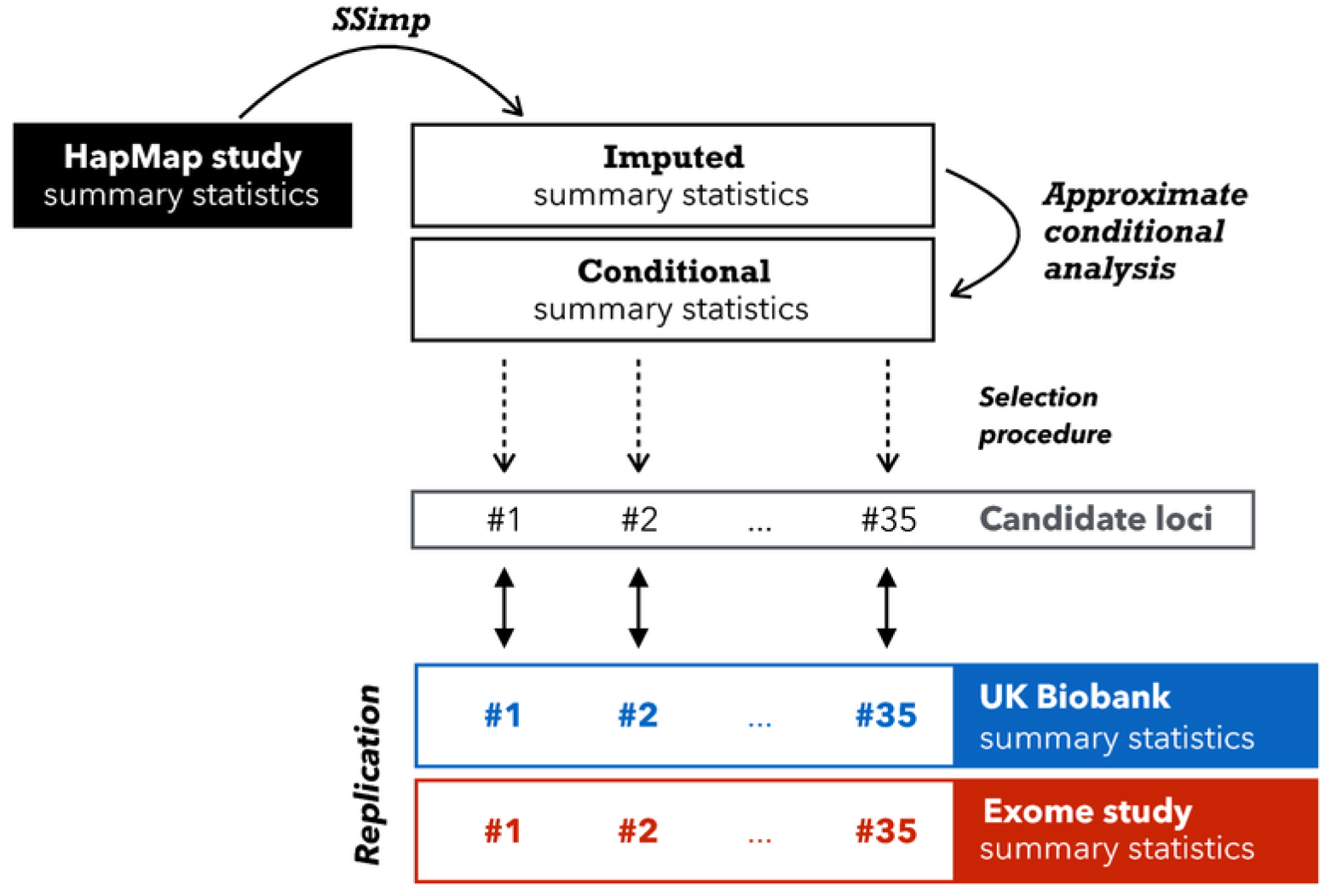
Overview of imputation and replication scheme. This illustration gives an overview how we used > 2M GIANT HapMap summary statistics (black rectangle) as tag SNVs to impute > 10M variants with MAF≥ 0.1% in UK10K. After adjusting the summary statistics for conditional analysis we applied a selection process that resulted in 35 candidate loci. To confirm these 35 loci we used summary statistics from UK Biobank (blue) as replication as well as summary statistics from the exome chip study, if available [13] (red). Loci that had not been discovered by the exome chip study, were termed *novel*.

We subjected all 9′276′018 variants with an imputation quality ≥ 0.3 to a scan for novel candidate loci. A region was defined as a candidate locus if at least one imputed variant was independent from any reported HapMap variant nearby (conditional *P*-value ≤ 10^−8^). We identified 35 such candidate loci. Within each locus we defined the imputed variant with the lowest conditional *P*-value as the top variant. All 35 variants are listed in S1 Table and locus-zoom plots are provided in S7 Fig.

Next, we used the UK Biobank to replicate the associations with height of these 35 candidate variants and subsequently grouped them into replicating (20 variants) and not replicating (15 variants) (at *α* = 0.05/35 level).

An overview of the 20 replicating variants is given in Table 2. One region had already been discovered in the GIANT exome chip study: **`rs28929474`**, located in gene *`SERPINA1`*. Fig 7 shows this region as locus-zoom plot with summary statistics from the HapMap study, *summary statistics imputation*, and the exome chip study. To annotate these 20 novel candidate variants further, we investigated whether they are eQTLs or associated with other traits. We report this in Table 3 where we list eQTLs detected by GTEx [18] and Table 4 that presents a curated association-trait list by Phenoscanner [19]. In the following we describe variants that replicated in UK Biobank which are either eQTLs or have previously been associated with another trait.

**Table 2.** Twenty replicating candidate loci for height.

| # | SNV | Chr | Pos | Allele<br>R/E | Gene<sup>(1)</sup> | MAF<br>(<sup>2</sup>) | SSimp<br>*P* | *N* | UK Biobank<br>*P* | *N* | Group |
| --- | --- | --- | --- | --- | --- | --- | --- | --- | --- | --- | --- |
| 1 | rs112635299<sup>(*)</sup> | 14 | 94838142 | G/T | - | 2.33% | 4.21E-14 | 234380 | 5.16E-77 | 336474 | (i) |
| 2 | rs76306191 | 1 | 155006451 | C/G | *DCST1* [E] | 20.30% | 6.51E-10 | 245908 | 2.74E-16 | 336474 | (ii) |
| 3 | rs73029259 | 6 | 164111348 | T/A | - | 12.77% | 7.61E-09 | 251161 | 1.02E-15 | 336474 | (ii) |
| 4 | rs67807996 | 1 | 149995265 | G/A | - | 40.16% | 1.48E-43 | 219605 | 2.75E-102 | 336474 | (iii) |
| 5 | rs12795957 | 11 | 67242216 | G/A | - | 5.46% | 1.52E-24 | 193457 | 1.75E-76 | 336474 | (iii) |
| 6 | rs503035 | 5 | 134353734 | A/G | - | 30.39% | 6.34E-24 | 248110 | 5.46E-39 | 336474 | (iii) |
| 7 | rs568777 | 6 | 81809121 | C/G | - | 26.61% | 7.08E-24 | 252456 | 3.11E-35 | 336474 | (iii) |
| 8 | rs75975831 | 19 | 17264961 | G/C | *MYO9B* [I] | 22.52% | 3.59E-10 | 233765 | 9.19E-22 | 336474 | (iii) |
| 9 | rs56006730 | 12 | 103132740 | G/A | - | 10.41% | 1.80E-09 | 250070 | 1.05E-19 | 336474 | (iii) |
| 10 | rs35374532 | 6 | 26163345 | A/AT | *HIST1H2BD* [I] | 38.85% | 2.97E-27 | 252327 | 8.64E-18 | 120086 | (iii) |
| 11 | rs80171383 | 11 | 46084677 | G/A | *PHF21A* [I] | 14.72% | 3.53E-16 | 247885 | 2.05E-16 | 336474 | (iii) |
| 12 | rs13108218 | 4 | 3443931 | A/G | *HGFAC* [I] | 39.72% | 2.15E-10 | 222502 | 5.05E-15 | 336474 | (iii) |
| 13 | rs428925 | 5 | 173022921 | G/A | - | 27.59% | 1.34E-16 | 206987 | 4.31E-13 | 336474 | (iii) |
| 14 | rs6085649 | 20 | 6665532 | A/G | - | 45.61% | 1.24E-09 | 251393 | 1.65E-12 | 336474 | (iii) |
| 15 | rs78566116 | 6 | 32396146 | G/T | - | 7.67% | 2.74E-19 | 248592 | 4.18E-12 | 336474 | (iii) |
| 16 | rs350889 | 19 | 4118481 | A/G | *MAP2K2* [I] | 24.28% | 8.17E-10 | 207571 | 7.11E-12 | 336474 | (iii) |
| 17 | rs7955819 | 12 | 20677958 | T/C | *PDE3A* [I] | 23.23% | 6.13E-10 | 250048 | 3.25E-08 | 336474 | (iii) |
| 18 | rs7971674 | 12 | 1513526 | A/T | *ERC1* [I] | 14.12% | 8.10E-09 | 240270 | 2.19E-07 | 336474 | (iii) |
| 19 | rs12939056 | 17 | 7754993 | G/A | *KDM6B* [E] | 43.26% | 1.09E-12 | 245015 | 7.64E-07 | 336474 | (iii) |
| 20 | rs58402222 | 1 | 46059835 | T/TA | *NASP* [I] | 45.72% | 7.50E-13 | 252901 | 1.79E-04 | 120086 | (iii) |

This table presents 20 regions that contain at least one imputed variant that is independent from top HapMap variants nearby and that replicated in the UK Biobank (at $\alpha = 0.05/35$ level). Each row represents one region (#), indicating the SNV with the lowest conditional *P*-value. The first seven columns provide general information for each variant, followed by the *P*-value and sample size from *summary statistics imputation*, *P*-value and sample size from the UK Biobank. The last column assigns each of the 35 candidate loci to one of three groups: candidate loci (i) that were reported by [13] already, (ii) that had no reported HapMap variant nearby and (iii) that had reported HapMap variants nearby. $\hat{r}_{\text{pred,adj}}^2$ of all variants listed was greater than or equal to 0.3. We provide a more detailed table for all 35 variants (both replicating and not replicating) in S1 Table.
(\*) rs28929474, exome chip study results: $P = 1.39 \times 10^{-45}$ , $N = 365'451$ .
(1) [I] intronic, [E] exonic, - intergenic.
(2) MAF was computed in UK10K.

**Fig 7.**
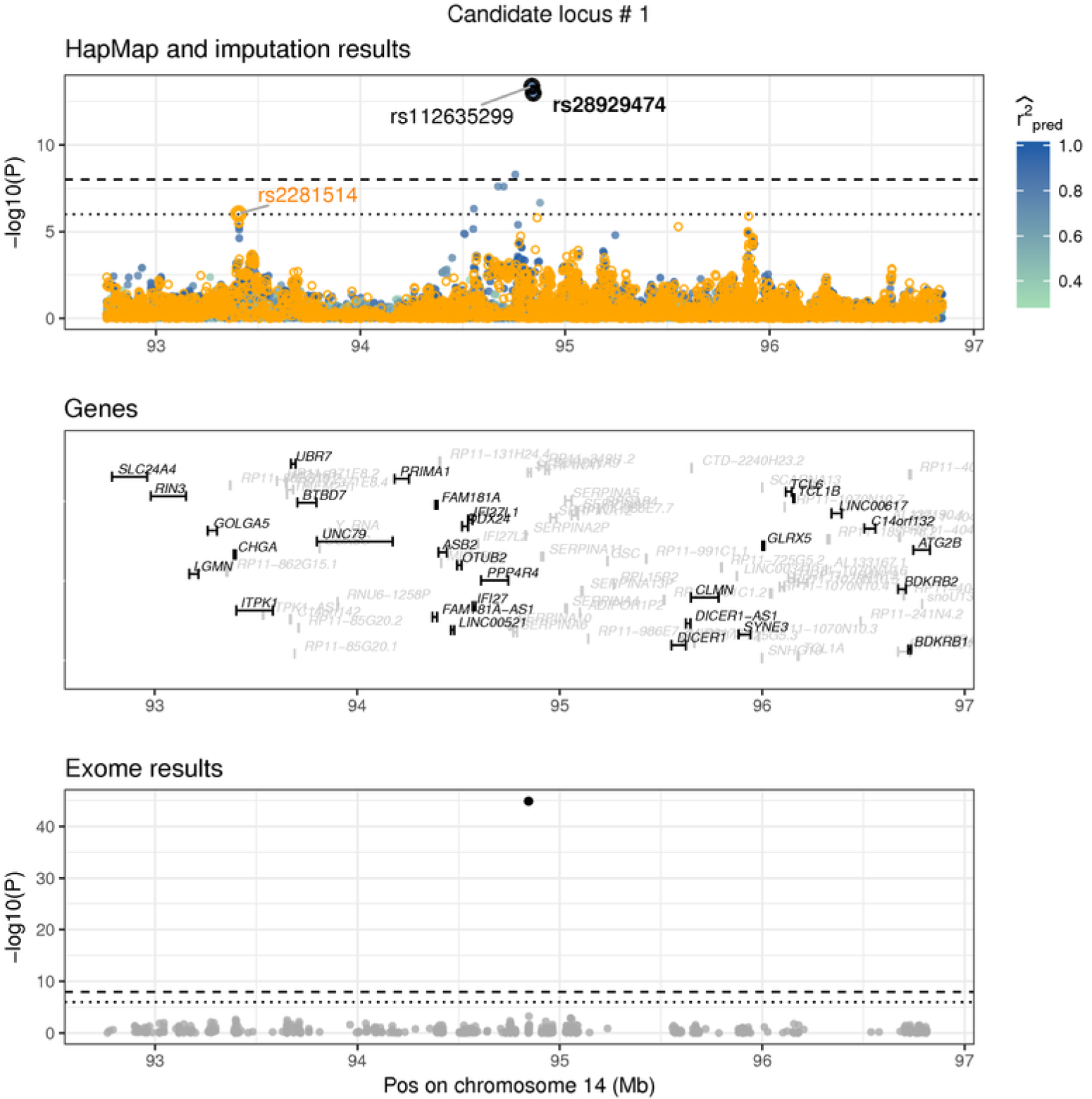
Replication of exome variant. **`rs28929474`** is a missense variant on chromosome 14 in gene SERPINA1, low-frequency (MAF=2.3%), imputed summary statistics (*P*_*SSimp*_ = 1:06×^−13^), replication in the UK Biobank (*P*_*UKBB*_ = 6:49×^−78^). **`rs112635299`** has the strongest signal in this region (*P* = 4:21 × 10^−14^), but is highly correlated to **`rs28929474`** (LD=0.95). This figure shows three datasets: Results from the HapMap and the exome chip study, and imputed summary statistics. The top window shows HapMap *P*-values as orange circles and the imputed *P*-values (using summary statistics imputation) as solid circles, with the colour representing the imputation quality (only 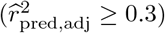 shown). The bottom window shows exome chip study results as solid, grey dots. Each dot represents the summary statistics of one variant. The x-axis shows the position (in Mb) on a ≥ 2 Mb range and the y-axis the −*log*10(*P*)-value. The horizontal line shows the *P*-value threshold of 10^−6^ (dotted) and 10^−8^ (dashed). Top and bottom window have annotated summary statistics: In the bottom window we mark dots as black if it is are part of the 122 reported hits of [13]. In the top window we mark the rs-id of variants that are part of the 122 reported variants of [13] in bold black, and if they are part of the 697 variants of [12] in bold orange font. Variants that are black (plain) are imputed variants (that had the lowest conditional *P*-value). Variants in orange (plain) are HapMap variants, but were not among the 697 reported hits. Each of the annotated variants is marked for clarity with a bold circle in the respective colour. The genes annotated in the middle window are printed in grey if the gene has a length < 50000 bp or is an unrecognised gene (**`RP−`**).

**Table 3.** GTEx annotation results for variants in eQTLs.

| # | SNV | $P_{SSimp}$ | $P_{UKBB}$ | GTEx tissue | Gene | $P$ |
| --- | --- | --- | --- | --- | --- | --- |
| 2 | rs76306191 | 6.51E-10 | 1.09E-07 | Artery_Tibial | *ZBTB7B* | 3.97E-09 |
|  |  |  |  | Thyroid | *DCST2* | 2.41E-08 |
| 5 | rs12795957 | 1.52E-24 | 6.17E-41 | Artery_Tibial | *RAD9A* | 6.48E-10 |
| 6 | rs503035 | 6.34E-24 | 8.06E-12 | Testis | *PITX1* | 2.91E-07 |
| 20 | rs58402222 | 7.50E-13 | 1.79E-04 | Cells_Transformed_fibroblasts | *MAST2* | 8.84E-23 |
|  |  |  |  | Cells_Transformed_fibroblasts | *CCDC163P* | 1.11E-19 |
|  |  |  |  | Cells_Transformed_fibroblasts | *TMEM69* | 2.16E-08 |
|  |  |  |  | Thyroid | *GPBP1L1* | 3.26E-11 |

This table shows SNVs which are significant eQTLs in GTEx [18]. We only report SNV-gene expression associations where the summary statistics pass the significance threshold of $\alpha = 10^{-6}$ . The first four columns represent the region number, SNV, $P$ -value from *summary statistics imputation* and the $P$ -value in the UK Biobank. The four remaining columns are information extracted from GTEx, with the tissue name, gene name, the $P$ -value of the association between the SNV and the gene expression, and the gene type. For each region, we only include the tissue with the lowest $P$ -value per SNV-gene associations. The full version of this table is available in S2 Table. # refers to the region number.

**Table 4.** Known trait association results for variants in Table 2.

| # | SNV | $P_{SSimp}$ | $P_{UKBB}$ | Study | PMID | Ancestry | Trait | $P$ | $N$ |
| --- | --- | --- | --- | --- | --- | --- | --- | --- | --- |
| 1 | rs112635299 | 4.21E-14 | 3.52E-25 | Wood | 23696881 | Mixed | Alpha 1 globulin | 2.51E-12 | 5278 |
|  |  |  |  | Kettunen J | 27005778 | European | Glycoprotein acetyls<br>mainly a1Lacid glycoprotein | 1.27E-10 | 17772 |
|  |  |  |  | Kettunen J | 27005778 | European | Total cholesterol in small LDL | 6.59E-10 | 20057 |
|  |  |  |  | Kettunen J | 27005778 | European | M.LDL.C | 4.03E-09 | 20060 |
|  |  |  |  | Kettunen J | 27005778 | European | Cholesterol esters in medium LDL | 6.19E-09 | 17774 |
|  |  |  |  | Kettunen J | 27005778 | European | Total lipids in medium LDL | 7.26E-09 | 17774 |
|  |  |  |  | Kettunen J | 27005778 | European | Total cholesterol in LDL | 8.66E-09 | 20060 |
|  |  |  |  | Kettunen J | 27005778 | European | Total lipids in small LDL | 1.56E-08 | 17774 |
|  |  |  |  | Kettunen J | 27005778 | European | Conc. of medium LDL particles | 1.67E-08 | 17774 |
|  |  |  |  | Kettunen J | 27005778 | European | Conc. of small LDL particles | 2.77E-07 | 17774 |
|  |  |  |  | Kettunen J | 27005778 | European | Cholesterol esters in large LDL | 4.72E-07 | 17774 |
|  |  |  |  | Kettunen J | 27005778 | European | Total cholesterol in large LDL | 7.36E-07 | 20053 |
|  |  |  |  | Kettunen J | 27005778 | European | Total lipids in large LDL | 9.86E-07 | 17774 |
| 15 | rs78566116 | 2.74E-19 | 9.80E-04 | Chen D | 21896673 | Mixed | HPV8 seropositivity in cancer | 3.30E-16 | 6885 |
|  |  |  |  | Okada Y | 24390342 | European | Rheumatoid arthritis | 3.80E-94 | 58284 |
|  |  |  |  | Okada Y | 24390342 | Mixed | Rheumatoid arthritis | 2.30E-90 | 80799 |
|  |  |  |  | IBDGC | 26192919 | European | Ulcerative colitis | 4.06E-08 | 27432 |

This table describes SNVs previously associated with other traits. The search was conducted with Phenoscanner [19]. We only list SNVs for which Phenoscanner had information available regarding GWAS traits or metabolites. The first four columns specify region, SNV-id, followed by the $P$ -value from *summary statistics imputation* and the $P$ -value from the UK Biobank. Column five to ten contain information extracted from Phenoscanner. We report the respective summary statistics that pass the significance threshold of $\alpha = 10^{-6}$ . # refers to the region number, conc. to concentration.

We can classify the 35 candidate loci into three categories (i), (ii) and (iii) that reflect the type of conditional analysis performed. Group (i) includes SNVs replicating already published exome chip associations (one locus), group (ii) includes SNVs that contain no reported HapMap variant nearby (three loci), and group (iii) includes SNVs that contain one or more reported independent HapMap variants nearby (31 loci). Replication success with UK Biobank is 1/1 in group (i), 2/3 in group (ii), 17/31 in group (iii). We only term categories (ii) and (iii) as *novel* candidate loci, therefore limiting the number of novel candidate loci to 34, with 19 replicating in UK Biobank.

Although group (ii) only contains loci that had no reported HapMap variants nearby, three candidate loci (#2, #3, #21 in S1 Table) contain borderline significant HapMap signals (*P*-value between 10^−6^ and 10^−8^ in [12]).

We observed that variants with higher MAF have higher chance to replicate. Among the 20 candidate variants that did replicate in UK Biobank, 19 were common and one a low-frequency variant (**`rs112635299`**, MAF = 2.32%). Conversely, among the 15 candidate variants that did not replicate in the UK Biobank, 10 are rare, three are low-frequency variants, and two are common.

**Locus #1:** rs112635299 (imputed *P*-value 4.21 x 10^−14^), is a proxy of rs28929474 (LD= 0.88), has been associated with alpha-1 globulin [20] and is associated with multiple lipid metabolites [21]. **`rs28929474`** was identified in the GIANT exome chip study to be height-associated (*P* = 1.39 × 10^−45^) [13]. The *P*-value calculated with *summary statistics imputation* was P = 1.06 × 10^−13^. **`rs28929474`** is a low-frequency variant (MAF= 2.3%) and replicates in the UK Biobank with *P* = 1.66 × 10 ^− 25^.

**Locus #2: `rs76306191`** is a common variant on chromosome 1, located in gene ***DCST1***. There was no reported HapMap variant nearby to condition on. However, the absolute correlation to the HapMap variant with the lowest *P*-value (> 10^−8^) in the same region was 0.8. One of the 122 variants reported by the exome chip study, **`rs141845046`**, was in this region, but had an imputed *P*-value > 10^−3^. **`rs76306191`** replicated in the UK Biobank with P = 1.09 x 10^−7^. rs76306191 is an eQTL in artery (tibial) for gene ***ZBTB7B*** and in thyroid gland for gene ***DCST2***.

**Locus #5: `rs12795957`** is a variant on chromosome 11 and an eQTL for gene ***RAD9A*** in artery (tibial).

**Locus #6: `rs503035`** is a variant on chromosome 5. It is an eQTL for gene ***PITX1*** in testis tissue. **`rs62623707`**, one of the 122 reported exome variants, was in this region, but had an imputed *P*-value > 10^−3^.

**Locus #15: `rs78566116`** is a variant on chromosome 6. **`rs78566116`** has been associated with HPV8 seropositivity in cancer [22], rheumatoid arthritis [23] and ulcerative colitis [24].

**Locus #20: `rs58402222`** is an intronic variant on chromosome 1, located in gene ***NASP***. It is an eQTL for genes ***CCDC163P***, ***MAST2*** and ***TMEM69*** in cells (transformed fibroblasts); and for ***GPBP1L1*** in thyroid tissue.

### Replication of 55/111 reported GIANT exome chip variants

Next, we focussed on 122 novel variants of Marouli *et al*. [13]. For this analysis we did not apply any MAF restrictions. Of these 122 variants, 11 variants were either not referenced in UK10K or on chromosome X, and were therefore not imputed, limiting the number of loci and variants to 111 (S3 Table). By grouping results below or above the *P*-value threshold of *α* = 0.05/111 we could classify variants into the ones that replicated and those that failed replication. This is summarised in Table 5 and S8 Fig, which shows that 55 of the 111 variants could be retrieved, four of them with MAF ≤ 5%.

Details to the imputation of all 111 variants are listed in S3 Table.

**Table 5.** 111 variants: Fraction of top variants in exome chip study retrieved with imputation of HapMap study.

| $\hat{r}_{\text{pred,adj}}^2$ | MAF | | |
| --- | --- | --- | --- |
|  | 5 – 50% | 1 – 5% | 0 – 1% |
| 0.7-1 | 65% (49/75) | 50% (4/8) | - |
| 0.3-0.7 | 67% (2/3) | 0% (0/17) | 0% (0/3) |
| 0-0.3 | - | - | 0% (0/5) |

This table presents *summary statistics imputation* results, limited to 111 variants identified as “novel” by [13]. We summarised the results according to their allele frequency and imputation quality category. For each subgroup we calculated the fraction of top exome variants that had a $P$ -value $\leq 0.05/111$ with *summary statistics imputation*.

## Discussion

In this article, we focussed on the comparison between *genotype* and *summary statistics imputation*. In contrast to previous work by others and us [10,11,25], here we systematically assessed the performance and limitations of *summary statistics imputation* through real data applications for different SNV subgroups characterised by allele frequency, imputation quality and association status (null/associated). In addition, we demonstrated the usefulness of *summary statistics imputation* to discover novel associated regions using existing association data. Note that in this paper we used an improved version of the original *summary statistics imputation* [11], which uses reference panel size dependent shrinking of the correlation matrix and incorporates variable sample size of tag SNVs.

Our study design has several limitations: for replication of summary statistics from European individuals we use the UK Biobank, which represents only a subset of all European ancestries and is genotype-imputed (instead of sequenced), but on the other hand provides a reliable resource due to its sample size. Furthermore, in UK Biobank, *genotype imputation* done for genotyped variants can only partially be compared to *genotype imputation* for untyped variants, as genotyped variants were used for phasing (therefore *genotype imputation* of genotyped variants is much easier and leads imputation qualities close to one, S4 Fig). Due to the small number of height-associated rare variants (13) we can not draw meaningful conclusions for this group and hence avoided their analysis.

The *summary statistics imputation* method itself has several limitations too. First, due to the size of publicly available sequenced reference panels we can not explore the performance of rare variants (MAF< 1%). Second, the imputation quality metric 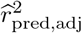 tends to be inaccurate in case of small reference panels. Third, the imputation of summary statistics of an untyped SNV is essentially the linear combination of the summary statistics of the tag SNVs (Eq. (1)). Such a model cannot capture non-linear dependence between tag- and target SNVs [9], which is often the case for rare variants [26,27]. In contrast, *genotype imputation* is able to capture such non-linear relationships by estimating the underlying haplotypes (a non-linear combination of tagging alleles). Furthermore, in case of *genotype imputation* it is sufficient that the relevant haplotypes are present in the reference panel, but the overall allele frequency does not need to match the GWAS allele frequency.

### Comparison of *summary statistics imputation* versus *genotype imputation*

We compared *summary statistics imputation* and *genotype imputation* by using individual-level data from the UK Biobank, where we evaluated the imputation results for 6′080 SNVs that were correlated with a height-associated variant (*associated* SNVs) and 31′567 that were not correlated to any height-associated SNVs on the same chromosome (*null* SNVs).

In general, imputation using *summary statistics imputation* leads to a larger RMSE than *genotype imputation* in all twelve SNV subgroups investigated (Fig 4). Among associated SNVs, *summary statistics imputation* performs similar to *genotype imputation* for well-imputed SNVs, but shows a trend for underestimation of the Z-statistics and lower correlation with the true effect size for medium- and badly-imputed SNVs (Fig 2). Conversely, *genotype imputation* has more consistent results for most of the twelve SNV subgroups (Fig 2 and 3), that is reflected in a correlation close to one between Z-statistics from genotype data and *genotype imputation* data.

### Underestimation for null and associated SNVs

Ultimately, the underestimation of imputed Z-statistics with *summary statistics imputation* leads to a lower type I error. We calculated power and FPR for both methods and observe that for a given significance threshold, *summary statistics imputation* has a lower FPR at the cost of lower power compared to *genotype imputation*. This effect is amplified for SNV groups with lower imputation quality 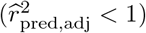. For associated SNVs with 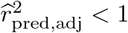 we expect an underestimation for associated SNVs due to the fact that we are imputing summary statistics under the null model, whereas for null SNVs with 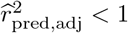 we expect an underestimation due to decreased variance of the *summary statistics imputation* estimation.

Ideally, for an unbiased estimation of causal and null SNVs, the imputed Z-statistics (Eq. (1)) should be divided by 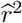. However, as the imputation quality 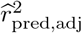 is noisily estimated from small reference panels (discussed below) and it is not guaranteed that the SNV we impute is causal, we risk to overestimate the summary statistics of associated SNVs. This is the reason why refrain from doing so.

S9 Fig shows the *P*-value distribution of *summary statistics imputation* for null SNVs with an accumulation of low −*log*10(*P*)-values for well-imputed SNVs and an accumulation of high −*log*10(*P*)-values for badly-imputed SNVs. We think that two factors are in play here. First, mostly due to polygenicity, the genomic lambda for height is λ_*GC*_ = 1.94, therefore we expect even seemingly null variants to show inflation. Second, for null SNVs, the sample variance of the imputed Z-statistics should be proportional to the average imputation quality. We calculated for each of the null SNV subgroups the ratio between the sample variance for Z-statistics from *summary statistics imputation* and the sample variance for Z-statistics from genotype data. For common null SNVs we observe a ratio that gradually decreases with imputation quality (0.89 for perfectly-, 0.79 for medium- and 0.68 for badly imputed SNVs). For low-frequency null variants the ratio is approximately 0.1 lower (0.82 for perfectly-, 0.70 for medium- and 0.52 for badly imputed SNVs). The inflation for well-imputed SNVs can be explained by the genomic lambda, while for badly-imputed SNVs it is aggravated by the underestimated standard error.

### Atypical allele frequency distribution and rare variants exclusion

Because the number of associated SNVs with MAF < 1% was too low (13 variants) to draw any meaningful conclusions, we refrained from analysing this MAF group. One other reason to exclude rare variants from this analysis is, that the reference panel used (UK10K) contains 3′871 individuals and therefore estimations for LD of rare variants are unreliable and rare variants can (in theory) only be covered down to MAF = 1/(2 · 3′871). We believe improving *summary statistics imputation* for rare variants will require not only larger reference panels to allow estimation of LD of rare variants, but also methods which would allow non-linear tagging of variants. It should be kept in mind that, just like for *genotype imputation*, even with very large reference panels, one will not be able to impute variants with extremely rare allele counts. To investigate these SNVs full genome sequencing is indispensable [28].

### Imputation quality

We find that our imputation quality measure 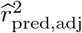 is conservative and probably underestimates the true imputation quality (S4 Fig). To calculate the imputation quality 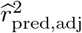, we need — similar to imputing summary statistics in Eq. (1) — to compute correlation matrices ***c*** and ***C*** estimated from a reference panel (Eq. (2)) and therefore encounter similar challenges as summary statistic imputation itself due to difficulties of reliable LD estimation.

The discrepancy in imputation quality metric between *summary statistics imputation* and *genotype imputation* (S4 Fig) can be explained by the fact that: (1) genotyped variants that were imputed too, were also used for phasing, (2) it is indeed more difficult to impute summary statistics using *summary statistics imputation*, and therefore the imputation quality is shifted towards zero, and (3) 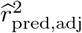 is an estimation that can either be erroneous due to choosing the wrong reference panel (and therefore 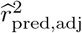 does not represent the true imputation quality) or it can be imprecise due to small sample size of the reference panel. For example, UK10K contains 3′871 individuals and is too small to precisely estimate these matrices (the standard error for a correlation estimated from *n* = 3′871 is 0.016), which becomes problematic in cases of low correlation.

### *Summary statistics imputation* of the height GWAS of the GIANT consortium

As a showcase of the utility of *summary statistics imputation* we imputed Wood *et al*. [12] to higher genomic resolution (limited to variants with MAF ≥ 0.1% as well as 111 previously reported exome variants) [13], then selected imputed variants that act independently from those reported in Wood *et al*. and replicated these with independent data.

While Wood *et al*. [12] is the largest height study to date in terms of number of markers (covering HapMap variants in 253′288 individuals), Marouli *et al*. [13] exceeds their sample size by more than 100′000 individuals, but is limited to 241′419 exome variants. The similarity between both GIANT studies made the exome chip study ideal for replication. We chose the UK Biobank as a second replication dataset, despite its limitation to individuals of British ancestry, as it covers more variants than the exome chip study.

The ultimate goal was to find new height-associated variants. To do so, we scanned through the imputed results and marked regions that had at least one variant confidently imputed 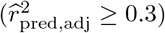 with an association with *P* ≤ 10^−8^ and that acted independently from any reported HapMap variant nearby. This search allowed us to identify 35 regions, of which one had already been identified in the recent GIANT height exome chip study (**`rs28929474`**) and 19 replicated in UK Biobank (at *α* = 0.05/35 level). Two candidate loci (#2, #3 in Table 2) that replicate in UK Biobank have borderline significant HapMap signals in close proximity (*P*-value between 10^−6^ and 10^−8^ in [12]) and were therefore not reported in the study in 2014.

The 15 non-replicating candidate loci were on average on a lower allele frequency spectrum (ten are rare, three are low-frequency variants, and two are common). Allele frequency was higher among the 20 replicating candidate variants (19 were common and one a low-frequency variant).

### Replicating GIANT exome chip imputation results

We then focussed on the *summary statistics imputation* of the the 111 reported exome chip variants [13]. Knowing from our previous findings that rare variants are challenging to impute due to reference panel size, we expected to retrieve a larger fraction of common and low-frequency than rare variants. S8 Fig shows that we retrieved 49.5% (55) of the variants when using a strict Bonferroni corrected threshold (at *α* = 0.05/111 level, Table 5). More specifically, of the 111 top variants (78 common, 25 low-frequency, eight variants rare) 83 variants were imputed with high confidence 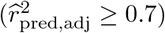. Of these, 53 were retrieved when using the typical candidate SNV threshold (0.05/111). Among variants with lower imputation quality only two common and medium-imputed variants could be retrieved. As shown in Fig 2 and 5, the power of *summary statistics imputation* decreases with lower MAF and imputation quality.

## Conclusion

In summary, we have evaluated the performance of our recently improved *summary statistics imputation* method in terms of different measures and shown that *summary statistics imputation* is a very efficient and fast method to separate null from associated SNVs. However, *genotype imputation* outperforms *summary statistics imputation* by a clear margin in terms of accuracy of effect size estimation. By imputing GIANT HapMap-based summary statistics we have demonstrated that *summary statistics imputation* is a rapid and cost-effective way to discover novel trait associated loci. We also highlight that the principal limitations of *summary statistics imputation* are rooted in the LD estimation and in imputing very rare variants with sufficient confidence.

## Materials and methods

### Comparison of *summary statistics imputation* versus *genotype imputation*

#### UK Biobank data

The UK Biobank [29] comprises health related information about 500′000 individuals based in the United Kingdom and aged between 40-69 years in 2006-2010. For our analysis we used Caucasians individuals (amongst people who self-identified as British) from the first release of the genetic data (*n* = 120′086). For SNVs, the number of individuals range between *n* = 3′431 and *n* = 120′082. Additionally to custom SNP array data, UK Biobank contains imputed genotypes [30]. A subset of 820′967 variants were genotyped and imputed, and 72*M* variants were imputed by UK Biobank, using **SHAPEIT2** and **IMPUTE2** [30].

#### Imputation of height GWAS summary statistics conducted in UK Biobank

We imputed GWAS Z-statistics (ran on directly genotyped data) within 1 Mb-wide regions, by blinding one at the time and therefore allowing the remaining SNVs to be used for tagging. As tag SNVs we used all SNVs except the focal SNV within a 1.5 Mb window.

#### Selection of regions and SNVs

We selected 706 regions in total, consisting of 535 loci containing height-associated SNVs [12,13] and 171 regions not containing any height-associated (all *P* ≥ 10^−5^) SNV. More specifically, within each height-associated region we only imputed SNVs that have LD_max_> 0.2. LD_max_ was defined as the largest squared correlation between a SNV and all height-associated SNVs on the same chromosome. In the 171 null regions we chose only those variants with LD_max_≤ 0.05 with any associated marker on the same chromosome. These selection criteria lead to 44′992 variants being imputed. We did not analyse palindromic SNVs (A/T and C/G) (3′306 variants), SNVs with missing genotypes for more than 36′024 (30%) individuals (2′317 variants), SNVs with MAF < 1% (3′010 variants). These restrictions left us with 37′467 of the 44′992 imputed SNVs.

#### Comparison of *summary statistics imputation* and *genotype imputation*

To compare the performance between *summary statistics imputation* and *genotype imputation* followed by association we compared each method to the directly genotyped data association as gold standard. We used RMSE, bias, correlation, and the regression slope (no intercept) to evaluate these approaches against the truth.

More precisely, the RMSE and the Bias for a set of *k* =1… *K* SNVs is:

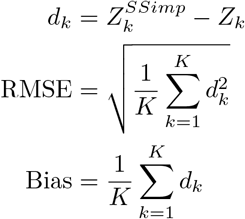

with 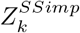 being the Z-statistic resulting from *summary statistics imputation* for SNV *k* and *Z*_*k*_ the Z-statistic resulting from genotype data for SNV *k* (our gold standard). Likewise, we replaced 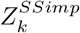 with 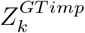, to calculate RMSE and bias for *genotype imputation*.

For genotype summary statistics from associated SNVs that resulted from data with partial sample size, we computed an upsampled Z-statistics, where *Z*_*u*_ represents the Z-statistics for SNV *u*, *N*_*u*_ the sample size of SNV *u* and *N*_max_ the maximal sample size within the study: 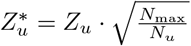. Whenever we use Z-statistics from associated genotype data we use this upsampled version ***Z***\*.

Additionally, we calculated power and false positive rate (FPR) for each method. For SNVs with a real association we calculated the power as the fraction of SNVs with a *P* ≤ *α*, whereas for SNVs with no association we calculated FPR as the fraction of SNVs with *P* ≤ *α*. We varied α between 0 and 1 and visualised FPR versus power for each method.

#### Stratifying results

The obtained (summary statistics) imputation results were grouped based on the imputed SNVs (i) being correlated (LD > 0.3) to any height-associated SNV on the same chromosome or being a null SNV (LD < 0.05); (ii) low-frequency (1% < MAF ≤ 5%) or common SNV (MAF > 5%); (iii) being badly-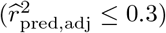, medium- 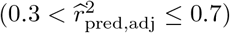 or well-imputed 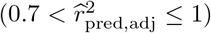. Height-associated SNVs are exclusively from 535 regions and termed *associated* SNVs, while SNVs not associated with height stem from 171 regions and are termed *null* SNVs. Throughout the manuscript, LD is estimated as the squared correlation [31].

### *Summary statistics imputation* of the height GWAS of the GIANT consortium

#### GIANT consortium summary statistics

In 2014 the GIANT consortium published meta-analysed height summary statistics involving 79 cohorts, 253′288 individuals of European ancestry, and 2′550′858 autosomal HapMap SNVs [12], leading to the discovery of 423 height-associated loci (697 variants). Later, Marouli *et al*. [13] published summary statistics of the exome array meta-analysis (241′419 SNVs in up to 381′625 individuals), finding 122 novel variants (located in 120 loci) associated with height. Of the 122 exome variants, four were not available in UK10K and seven were on chromosome X, and could therefore not be imputed (because [12] did not include chromosome X), leaving 111 variants. We refer to the summary statistics by Wood *et al*. [12] as HapMap study, and to Marouli *et al*. [13] as exome chip study.

#### *Summary statistics imputation* of Wood *et al.*

We imputed all non-HapMap variants that were available in UK10K, using the summary statistics in [12] as tag SNVs. In general, we only imputed variants with MAF_UK10K_ ≥ 0.1% (this allows a minimal allele count of 8 ≃ 0.001 · 3781 · 2), except for the 111 exome variants reported in [13], which we imputed regardless of their MAF. We divided the genome into 2′789 core windows of 1 Mb. We imputed the summary statistics of each variant using the tag SNVs within its respective window and 250 Kb on each side. Fig 6 gives an overview of the datasets and methods involved.

#### Definition of a candidate locus

After applying *summary statistics imputation* we screened for SNVs with 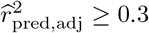 and an (imputed) *P*-value ≤ 10^−8^ and applied conditional analysis, aiming to limit the results to SNVs acting independently from known HapMap findings. The significance threshold of 10^−8^ was chosen based on the effective number of SNVs evaluated (< 9′276′018). For each imputed 1 Mb window, we started the conditional analysis by defining two sets of SNVs. The first set contained all imputed SNVs that had an imputed *P*-value ≤ 10^−8^, ranging from position bp^(1)^ to bp^(2)^. The second SNV set contained all reported HapMap SNVs (697 in total) within a range of bp^(1)^ — 1 Mb and bp^(2)^ + 1 Mb. Having two SNV sets - the first set with newly detected variants, the second set with reported HapMap variants - we could then condition each SNV in the first set on all SNVs in the second set, using approximate conditional analysis [32] and UK10K as the reference panel. Next, we declared a region as a candidate locus if at least one imputed variant in that locus had a conditional *P*-value ≤ 10^−8^. Finally, we performed a conditional analysis for nearby candidate loci (neighbouring windows), to avoid double counting. In each candidate locus we report the imputed variant with the smallest conditional *P*-value as the top variant.

#### Replication of candidate loci emerging from *summary statistics imputation*

We replicate our findings using our UK Biobank height GWAS results and for SNVs present on the exome chip we also use the recent height GWAS [13]. For both attempts to replicate our findings, UK Biobank and the exome chip study, the significance threshold for replication is *α* = 0.05/*k*, with *k* as the number of candidate loci.

For replication using UK Biobank we used summary statistics based on the latest release of genetic data with *n* = 336′474 individuals, provided by the Neale lab [33]. For SNVs that were not present in the latest release we used summary statistics from the first release of genetic data (*n* = 120′086)).

#### Annotation of candidate loci

We use two databases to annotate newly discovered SNVs. First, we use GTEx [18], an eQTL database with SNV-gene expression association summary statistics for 53 tissues. Second, we conduct a search in Phenoscanner [19], to identify previous studies (GWAS and metabolites) where the newly discovered SNVs had already appeared. For these two databases we report the respective summary statistics that pass the significance threshold of *α* = 10^−6^. We only extract the information for variants that were defined as as novel discoveries.

#### Reference panels

To estimate LD structure in *C* and *c* (Eq. (1)) we used 3′781 individuals from UK10K data [34,35], a reference panel of British ancestry that combines the TWINSUK and ALSPAC cohorts.

#### Software

All analysis was performed with **R-3.2.5** [36] programming language, except GWAS summary statistics computation for UK Biobank genotype and genotype imputed data, for which **SNPTEST-5.2** [37] was used. For summary statistics imputation we used **SSIMP** [38].

## Supporting information

**S1 Fig. UK Biobank: Absolute frequencies of allele frequency and imputation quality of imputed SNVs.** This figure shows how many of the null and associated SNVs were categorised into common, low-frequency and rare MAF subgroups, and into well-imputed, medium imputed and badly imputed imputation subgroups. Associated SNVs are presented in the left window, and null SNVs are presented in the right window. MAF category (x-axis), # of SNVs on the y-axis, colour refers to imputation quality category.

*Link to S1 Fig*.

**S2 Fig. UK Biobank: Relative frequencies of imputation quality within each allele frequency group.** This figure shows the fraction of badly-, medium- and well-imputed SNVs within each MAF subgroup. Null and associated SNVs were categorised into common, low-frequency and rare MAF subgroup, and into well-imputed, medium imputed and badly imputed imputation subgroup. Associated SNVs are presented in the left window, and null SNVs are presented in the right window. MAF category (x-axis), fraction of SNVs on the y-axis, colour refers to imputation quality category. Numbers within the stacked barplot refer to the number of SNVs imputed in each subgroup.

*Link to S2 Fig*.

**S3 Fig. UK Biobank: Comparison of imputation quality methods. MACH** 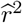 [39] (x-axis) versus IMPUTE's info measure used by *genotype imputation* (y-axis). To avoid clumping of dots, we used tiles varying from grey (few dots) to black (many dots). The identity line is dotted.

*Link to S3 Fig*.

**S4 Fig. UK Biobank: Comparison of imputation quality methods.**

**IMPUTE**’s info measure used by *genotype imputation* (x-axis) vs 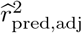 used by *summary statistics imputation* (y-axis). To avoid clumping of dots, we used tiles varying from grey (few dots) to black (many dots). The identity line is dotted.

*Link to S4 Fig*.

**S5 Fig. UK Biobank: FPR versus power.** This figure compares the false positive rate (FPR) (x-axis) versus the power (y-axis) for *genotype imputation* (blue) and *summary statistics imputation* (green) for different significance thresholds (*α*). The coloured dots represent the left-top-most point, with the printed α next to it. Results are grouped according to MAF (columns) and imputation quality (rows) categories. A zoom into the area of FPR between 0 and 0.1 can be found in Fig 5.

*Link to S5 Fig*.

**S6 Fig. GIANT: Concordance between genotyping and exome chip results**

This graph shows the Z-statistics of the exome chip study on the x-axis versus the Z-statistics of SNP-array study on the y-axis. Each dot shows one of the 2′601 variants that had LD_max_ > 0.1 (LD with one of the top variants in the exome [13] or HapMap study [12]). To make the density more visible, dots have been made transparent. The solid line indicates a linear regression fit, with the slope in the top right corner (including the 95%-confidence interval in brackets). The dashed line represents the ratio between the two median sample sizes 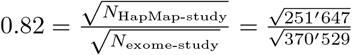

*Link to S6 Fig*.

**S7 Fig. Locus-zoom plots of all 35 regions. Filename according to column ‘filename’ S1 Table**
This figure shows three datasets: Results from the HapMap and the exome chip study, and imputed summary statistics. The top window shows HapMap *P*-values as orange circles and the imputed *P*-values (using *summary statistics imputation*) as solid circles, with the colour representing the imputation quality (only 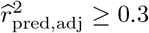 shown). The bottom window shows exome chip study results as solid, grey dots. Each dot represents the summary statistics of one variant. The x-axis shows the position (in Mb) on a ≥ 2 Mb range and the y-axis the −*log*10(*P*)-value. The horizontal line shows the *P*-value threshold of 10^−6^ (dotted) and 10^−8^ (dashed). Top and bottom window have annotated summary statistics: In the bottom window we mark dots as black if it is are part of the 122 reported hits of [13]. In the top window we mark the rs-id of variants that are part of the 122 reported variants of [13] in bold black, and if they are part of the 697 variants of [12] in bold orange font. Variants that are black (plain) are imputed variants (that had the lowest conditional *P*-value). Variants in orange (plain) are HapMap variants, but were not among the 697 reported hits. Each of the annotated variants is marked for clarity with a bold circle in the respective colour. The genes annotated in the middle window are printed in grey if the gene has a length < 5′000 bp or is an unrecognised gene (**`RP−`**).

*Link to S7 Fig*.

S8 Fig. Summary of exome results replication
This graph shows for all 111 variants the −*log*10(*p*)-value of the exome chip study on the x-axis and the imputed −*log*10(*p*)-value on the y-axis. The first row refers to the highest imputation quality (between 0.7 and 1), with the columns as the different allele frequency categories. The number of dots in each window is marked top left. The vertical and horizontal dotted lines mark the significance threshold of −*log*10(0.05/111) (dashed). The width of the x-axis is proportional to the range of the y-axis. For MAF and 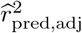 notation, the lower bound is excluded while the upper bound is included. For example, 1 — 5% is equivalent to 1 < MAF ≤ 5. *Link to S8 Fig*.

**S9 Fig. UK Biobank: Distribution of *P*-values from *summary statistics imputation*.** These QQ-plots show the distribution of *p*-values resulting from *summary statistics imputation*, for associated variants (left window), null variants (right window). The colours refer to the imputation quality categories. Note that the *P*-value in these plots are not λ_*GC*_ corrected.

*Link to S9 Fig*.

**S1 Table. GIANT: Detailed results of 35 candidate loci.** This table presents details of the 35 candidate loci discovered with *summary statistics imputation*. Within each candidate locus, we provide for the top variant the imputation results (**`.imp`**), along with conditional analysis results (**`.cond`**), the UK Biobank replication (**`.ukbb`**, whether it replicated or not (**`replication`**), and (if available) the exome chip study results (**`.exome`**). **`filename`** shows the filename of the locus-zoom plot in S7 Fig. **`SNP.cond.info`** presents each HapMap SNV used for conditional analysis, including its MAF, LD between the HapMap SNV and the imputed SNV, and a reversed conditional analysis result (HapMap variant conditioned on the imputed SNV). The column **`Group`** classifies each row into candidate loci (i) that were reported by [13] already, (ii) that had no reported HapMap variant nearby, (iii) that had at least one reported HapMap variants nearby. P = *P*-value, N = sample size, r2 = imputation quality, eff = effect size, EAF = effect allele frequency, MAF = minor allele frequency. If a candidate locus was not available in the UK Biobank, we provide a replication for a second variant that is in high LD with the primary variant, hence duplicated region numbers for some candidate loci.

*Link to S1 Table*.

**S2 Table. GTEx annotation results for variants in eQTLs**
This table shows SNVs which are significant eQTLs in GTEx [18]. We only report SNV-gene expression associations where the summary statistics pass the significance threshold of *α* = 10^−6^. The first four columns represent the region number, SNV, *P*-value from *summary statistics imputation* and the *P*-value in the UK Biobank. The three remaining columns are information extracted from GTEx, with the tissue name, gene name and the *P*-value of the association between the SNV and the gene expression. For each region, we order SNV-gene-tissue associations according to their *P*-value. # refers to the region number. *Link to S2 Table*.

**S3 Table. GIANT: Results of 122 exome variants.** This table presents the summary statistics imputation results (**`.imp`**) for all 122 variants shown as “novel” in [13]. The right hand part of the table shows the original exome chip results for comparison (**`.exome`**). P = *P*-value, N = sample size, r2 = imputation quality, eff = effect size, EAF = effect allele frequency. 11 variants were not referenced in UK10K or on chromosome X and therefore not imputed (see column ‘comment’). The position corresponds to hg19.

*Link to S3 Table*.

## Acknowledgments

This research has been conducted using the UK Biobank Resource. Eleonora Porcu, Jonathan Sulc, Kaido Lepik and Ninon Mounier gave valuable comments on an earlier draft of the manuscript.

## References

1. 1000 Genomes Project Consortium. A map of human genome variation from population-scale sequencing. Nature. 2010;467(7319):1061–1073.

2. Haplotype Reference Consortium. A reference panel of 64,976 haplotypes for genotype imputation. Nature genetics. 2016;48(10).

3. Howie B, Marchini J, Stephens M. Genotype Imputation with Thousands of Genomes. G3. 2011;1(6):457–470.

4. Fuchsberger C, Abecasis GR, Hinds DA. Minimac2: Faster genotype imputation. Bioinformatics. 2015;31(5):782–784.

5. Burgess S, Butterworth A, Thompson SG. Mendelian randomization analysis with multiple genetic variants using summarized data. Genetic Epidemiology. 2013;37(7):658–665.

6. Yang J, Benyamin B, McEvoy BP, Gordon S, Henders AK, et al. Common SNPs explain a large proportion of the heritability for human height. Nat Gen. 2010;42(7):565–569.

7. Bulik-Sullivan BK, Loh PR, Finucane HK, Ripke S, Yang J, Patterson N, et al. LD Score regression distinguishes confounding from polygenicity in genome-wide association studies. Nature Genetics. 2015;47(3):291–295.

8. Pickrell JK. Joint analysis of functional genomic data and genome-wide association studies of 18 human traits. 2014;94(4):559–573.

9. Pasaniuc B, Price AL. Dissecting the genetics of complex traits using summary association statistics. Nat Rev Genet. 2017;18(2):117–127.

10. Rüeger S, McDaid A, Kutalik Z. Improved imputation of summary statistics for realistic settings. bioRxiv. 2017;doi:10.1101/203927.

11. Pasaniuc B, Zaitlen N, Shi H, Bhatia G, Gusev A, Pickrell J, et al. Fast and accurate imputation of summary statistics enhances evidence of functional enrichment. Bioinformatics. 2014;30(20).

12. Wood AR, et al. Defining the role of common variation in the genomic and biological architecture of adult human height. Nature Genetics. 2014;46(11).

13. Marouli E, et al. Rare and low-frequency coding variants alter human adult height. Nature. 2017;.

14. Schäfer J, Strimmer K. A shrinkage approach to large-scale covariance matrix estimation and implications for functional genomics. Statistical applications in genetics and molecular biology. 2005;4.

15. Lee D, Williamson VS, Bigdeli TB, Riley BP, Fanous aH, Vladimirov VI, et al. JEPEG: a summary statistics based tool for gene-level joint testing of functional variants. Bioinformatics. 2014;31(8).

16. Gao X, Starmer ÃJ, Martin ER. A Multiple Testing Correction Method for Genetic Association Studies Using Correlated Single Nucleotide Polymorphisms. Genetic Epidemiology. 2008;369:361–369.

17. Lango Allen H, et al. Hundreds of variants clustered in genomic loci and biological pathways affect human height. Nature. 2010;467(7317):832–838.

18. Ardlie KG, et al. The Genotype-Tissue Expression (GTEx) pilot analysis: Multitissue gene regulation in humans. Science. 2015;348(6235):648–660.

19. Staley JR, Blackshaw J, Kamat MA, Ellis S, Surendran P, Sun BB, et al. PhenoScanner: a database of human genotype-phenotype associations. Bioinformatics. 2016;32(20):3207.

20. Wood AR, Perry JRB, Tanaka T, Hernandez DG, Zheng HF, Melzer D, et al. Imputation of Variants from the 1000 Genomes Project Modestly Improves Known Associations and Can Identify Low-frequency Variant - Phenotype Associations Undetected by HapMap Based Imputation. PLOS ONE. 2013;8(5):1–13.

21. Kettunen J, Demirkan A, Würtz P, Draisma HHMM, Haller T, Rawal R, et al. Genome-wide study for circulating metabolites identifies 62 loci and reveals novel systemic effects of LPA. Nature Communications. 2016;7:11122.

22. Chen D, McKay JD, Clifford G, Gaborieau V, Chabrier A, Waterboer T, et al. Genome-wide association study of HPV seropositivity. Human Molecular uGenetics. 2011;20(23):4714–4723.

23. Okada Y, Wu D, Trynka G, Raj T, Terao C, Ikari K, et al. Genetics of rheumatoid arthritis contributes to biology and drug discovery. Nature. 2014;506(7488):376–381.

24. Liu JZ, van Sommeren S, Huang H, Ng SC, Alberts R, Takahashi A, et al. Association analyses identify 38 susceptibility loci for inflammatory bowel disease and highlight shared genetic risk across populations. Nature Genetics. 2015;47(9):979–989.

25. Lee D, Bigdeli TB, Riley BP, Fanous AH, Bacanu SA. DIST: Direct imputation of summary statistics for unmeasured SNPs. Bioinformatics. 2013;29(22).

26. Wood AR, Tuke MA, Nalls MA, Hernandez DG, Bandinelli S, Singleton AB, et al. Another explanation for apparent epistasis. Nature. 2014;514(7520):E3–E5.

27. Hemani G, Shakhbazov K, Westra Hj, Esko T, Henders AK, Mcrae AF, et al. transcription in humans. Nature. 2014;508(7495):249–253.

28. Wu Y, Zheng Z, Visscher PM, Yang J. Quantifying the mapping precision of genome-wide association studies using whole-genome sequencing data. Genome Biology. 2017;18(1):86.

29. Sudlow C, Gallacher J, Allen N, Beral V, Burton P, Danesh J, et al. UK Biobank: An Open Access Resource for Identifying the Causes of a Wide Range of Complex Diseases of Middle and Old Age. PLoS Medicine. 2015;12(3):1–10.

30. UK Biobank Phasing and Imputation Documentation; 2015. https://biobank.ctsu.ox.ac.uk/crystal/docs/impute_ukb_v1.pdf.

31. Slatkin M. Linkage disequilibrium-understanding the evolutionary past and mapping the medical future. Nature reviews Genetics. 2008;9(6):477–85.

32. Yang J, Ferreira T, Morris AP, Medland SE, Madden PaF, Heath AC, et al. Conditional and joint multiple-SNP analysis of GWAS summary statistics identifies additional variants influencing complex traits. Nature Genetics. 2012;44(4):369–375.

33. Abbott L, Anttila V, Aragam K, Bloom J, Bryant S, Churchhouse C, et al. Rapid GWAS of thousands of phenotypes for 337′000 samples in the UK Biobank; 2017. Available from: http://www.nealelab.is/blog/2017/7/19/rapid-gwas-of-thousands-of-phenotypes-for-337000-samples-in-the-uk-biobank.

34. Moayyeri A, Hammond CJ, Valdes AM, Spector TD. Cohort profile: Twinsuk and healthy ageing twin study. International Journal of Epidemiology. 2013;42(1):76–85.

35. Boyd A, Golding J, Macleod J, Lawlor DA, Fraser A, Henderson J, et al. Cohort profile: The ‘Children of the 90s’-The index offspring of the avon longitudinal study of parents and children. International Journal of Epidemiology. 2013;42(1):111–127.

36. R Core Team. R: A Language and Environment for Statistical Computing; 2015. Available from: http://www.R-project.org/.

37. Marchini J, Howie B, Myers S, McVean G, Donnelly P. A new multipoint method for genome-wide association studies by imputation of genotypes. Nature genetics. 2007;39(7):906–13.

38. McDaid A, Rüeger S, Kutalik Z. SSIMP: Summary statistics imputation software; 2017. http://wp.unil.ch/sgg/summary-statistic-imputation-software/.

39. Li Y, Willer CJ, Ding J, Scheet P, Abecasis GR. MaCH: Using sequence and genotype data to estimate haplotypes and unobserved genotypes. Genetic Epidemiology. 2010;34(8):816–834.

